# Genomic Analysis Reveals Recent Population Decline and Exceptionally Low Genome-Wide Heterozygosity of the Critically Endangered Philippine Eagle, *Pithecophaga jefferyi* (Aves: Accipitridae)

**DOI:** 10.1101/2025.10.27.684716

**Authors:** Dhan Mikhail Perdon, Franchesca Pascual, Francis Tablizo, Carlo Lapid, John Michael Egana, Renato Jacinto Mantaring, Kris Punayan, Shiela Mae Araiza, Jo-Hannah Llames, Ma. Celeste Abad, Juan Carlos Gonzalez, Jayson Ibañez, Cynthia Palmes Saloma

## Abstract

The Philippine Eagle (*Pithecophaga jefferyi*), is one of the rarest eagles in the world and is the national bird of the Philippines. It is listed by the International Union for Conservation of Nature (IUCN) as a critically endangered raptor and has been the subject of an intensive captive breeding program which started more than 30 years ago to ensure the survival and conservation of the species. To infer the genetic diversity and demographic history of the Philippine Eagle, we sequenced and analyzed the genomes of 35 individuals under the captive breeding program of the Philippine Eagle Foundation. Phylogenetic analysis of the draft reference sequence generated in this study placed *P. jefferyi* within the subfamily Circaetinae of Accipitridae. Demographic history reconstruction from genome-wide variants revealed two historical bottlenecks, as well as an ongoing population decline which was found to predate documented deforestation in the Philippines. This observation suggests that unobserved natural or anthropogenic factors might have severely affected the Philippine Eagle population long before habitat fragmentation. Genome-wide heterozygosity estimates placed the Philippine Eagle as having one of the lowest genome-wide heterozygosity levels measured among raptors. This highlights the precarious genetic state of the Philippine Eagle, as critically low heterozygosity raises risks of inbreeding depression, reduced reproductive success, and increased vulnerability to diseases, climate change, and habitat loss. The genomic resources generated in this study can therefore guide conservation strategies such as breeding program design, genetic monitoring, and other efforts to diversify existing populations to ensure both demographic stability and genetic resilience of the Philippine Eagle.

**Subject Areas:** Philippine Eagle, Genomics, Genome-wide heterozygosity, Demographic History, Conservation, Biodiversity, Bioinformatics

## Introduction

The Philippine Eagle (*Pithecophaga jefferyi*) is one of the largest and most powerful forest raptors [1], and is officially recognized as the national bird of the Philippines [2]. It is found across four islands: Mindanao, Leyte, Samar, and Luzon, though its population is now fragmented and scattered across montane and lowland tropical forests [3, 4]. As forest-dependent raptors needing large land areas for hunting territory, it is susceptible to habitat destruction and fragmentation [5, 6], with many planned mining concessions overlapping spatially with high-suitability nesting areas [4, 7, 8]. Because of this, the Philippine Eagle is classified as ‘Critically Endangered’ on the IUCN Red List (also listed as CR under the Philippine Red List DAO 2019-09) and is one of the most threatened birds globally [9, 10]. Its population has drastically declined over the past five decades, primarily due to habitat loss from deforestation, agricultural expansion, and mining, as well as persecution. Mindanao Island remains the species’ stronghold, holding the largest population [7, 9]. Recent estimates, derived from Species Distribution Models (SDMs) and home-range data, suggest a global population of approximately 392 breeding pairs, or 784 mature individuals [4], emphasizing the urgent need for comprehensive conservation strategies.

Much like other imperiled raptors in Mindanao, the Philippine Eagle acts as a sentinel species, whose population status can indicate ecosystem health. Additionally, targeting conservation actions to the raptor could benefit a wide array of co-occurring taxa as an umbrella species [11, 12, 13]. Despite its ecological significance, broader biological investigations on its molecular information have been limited. Previous studies have investigated haplotype and nucleotide diversity of the mitochondrial control region of captive Philippine Eagles [14], and more recent studies assembled their partial mitogenomes and conducted nucleotide diversity analysis and k-means clustering to elucidate population structure within Mindanao [15].

While these studies have largely focused on the mitochondrial genome due to its cost-effective sequencing and wide use as a biomarker, mitochondrial data represent only a small fraction of the Philippine Eagle’s genetic information and may not fully capture patterns of variation within the whole nuclear genome [16, 17, 18]. Although a previous genome for the Philippine Eagle has been published in NCBI [19], the resulting assembly was assessed and found to contain a high proportion of ambiguous bases and unresolved regions.

The lack of a comprehensive and highly-documented genomic resource has previously hindered detailed genetic analyses, preventing a deeper understanding of the species’ population biology, crucial for effective conservation management. Similar to local initiatives that have demonstrated the value of genomic resources for other endemic and endangered species in the Philippines [20], a comprehensive genomic resource for the Philippine Eagle is vital for addressing its significant conservation challenges. This paper addresses this critical gap by presenting a draft genome assembly of the Philippine Eagle, which aims to support ongoing evolutionary research and conservation efforts. Analysis of genome assembly quality, BUSCO completeness, and comparative genomics to closely-related raptors [21] has been performed to assess quality, completeness, and accuracy of the contig-level assembly.

Initial analyses leveraging this novel genomic dataset, such as the estimation of genome-wide heterozygosity and reconstruction of demographic history, crucial for guiding conservation actions, were also performed [22]. This dataset represents the initial phase of a broader effort to generate the necessary resources for the integration of genomic information in conservation and breeding programs [23]. For future studies, this genomic resource can serve as a reference for designing molecular markers that can inform long-term monitoring and population management of the Philippine Eagle [9, 11] and support evidence-based strategies for conservation of the eagle and other co-occurring endemic species [11].

## Methods

### Sample collection and DNA extraction

Blood samples were collected from thirty-five Philippine Eagles at the Philippine Eagle Center of the Philippine Eagle Foundation in Davao City, Philippines. Samples were transported under the Wildlife Transport Permit No. DVO-WLTP-202400251 from the Department of Environment and Natural Resources (DENR). The samples were collected in K2 EDTA tubes and stored at 4°C before DNA extraction using the DNeasy Blood and Tissue Kit (Qiagen). DNA quality and quantity were assessed using agarose gel electrophoresis (AGE), spectrophotometry (NanoDrop, ThermoFisher Scientific), and fluorometry (Qubit 2.0 Fluorometer, Life Technologies). Extracted DNA was stored at −20°C until further processing.

### Library preparation and sequencing

Genomic libraries were constructed from 100 ng of high-quality genomic DNA using the MGIEasy FS DNA Library Prep Kit (MGI). Library concentrations were quantified using the Qubit dsDNA Broad Range Assay (Life Technologies), and average fragment sizes were determined using the Tapestation 2000 (Agilent Technologies). Libraries were prepared for DNA Nanoball sequencing using the MGIEasy Circularization Module (MGI) and sequenced on the DNBSEQ-G400 platform, generating 150-bp paired-end reads with approximately 30x genome coverage per individual.

### Genome assembly and quality assessment

Raw sequencing reads for each Philippine Eagle sample were quality-trimmed using FASTP v1.0.1, and individual genomes were assembled *de novo* using SOAP denovo2 v2.4.2 with k-mer sizes of 61, 71, 81, and 91 [24, 25]. Assembly quality was evaluated using QUAST v5.2.0, with the k-mer 71 assembly showing optimal assembly metrics [26]. To further improve assembly quality, individual assemblies were concatenated into a single FASTA file to simulate long reads, which were then re-assembled using long-read assemblers Flye v2.9.1 and Redbean v2.5.0 to generate higher-quality “meta-assemblies” [27, 28]. Assembly completeness was assessed using BUSCO v5.7.0 with aves_odb10 [29]. While Redbean produced superior QUAST metrics, Flye identified more complete BUSCO genes. Consequently, the Flye meta-assembly was selected for downstream analyses based on better gene completeness.

### Genome annotation

Transfer RNAs and ribosomal RNAs were predicted using tRNAscan-SE v2.0.12 and barrnap v0.9, respectively [30, 31]. Repetitive elements were identified and subsequently masked using RepeatMasker v4.1.6 [32]. To exclude sex-linked sequences, the repeat-masked genome to the Golden Eagle reference assembly (*Aquila chrysaetos*, GCF_900496995.4) using minimap v2.28, enabling identification of contigs associated with the Z and W chromosomes [33]. Only autosomal contigs were retained for gene annotation to avoid any errors introduced by the sex chromosomes [34].

A comprehensive gene annotation strategy was employed, integrating multiple prediction approaches through EVidenceModeler (EVM) v2.1.0 [35]. Homology-based prediction with GeMoMa v1.9 and protein sequences from three closely related Accipitridae species: golden eagle (*Aquila chrysaetos*, GCF_900496995.4), white-tailed eagle (*Haliaeetus albicilla*, GCF_947461875.1), and harpy eagle (*Harpia harpyja*, GCA_026419925.1) [36]. For ab initio prediction, two AUGUSTUS v3.5.0 models were trained and applied: one using raptor sequences (derived from GeMoMa output) and another using chicken (*Gallus gallus*) sequences [37]. The autoAug workflow was utilized, which encompassed custom training set generation (autoAug.pl), parameter optimization (autoAugTrain.pl), and genome-wide gene prediction (autoAugPred.pl). Broad protein evidence was elucidated through miniprot v0.18 alignment of avian proteins (taxonomy ID 8782) from the UniProt database [38]. The initial dataset was filtered using seqkit v2.10.0 to remove fragmented, partial, or exceptionally long sequences (>5,000 amino acids), reducing the dataset to 5,728,600 proteins [39]. Further clustering using CD-HIT v4.6.8 at 90% sequence identity yielded 965,952 bird protein sequences for alignment [40].

All annotation outputs were first converted to EVM-compatible formats using EvmUtils conversion scripts. Final consensus gene models were predicted using EVM with the following weights: AUGUSTUS-raptor = 5, GeMoMa = 4, miniprot = 2, AUGUSTUS-chicken = 1. Annotation quality of all annotation approaches was evaluated using BUSCO in protein mode.

Functional annotation was conducted using eggNOG-mapper v2 to assign Gene Ontology (GO) terms and orthologous groups [41]. The same approach was applied to the three reference Accipitridae species for comparative analysis. GO term distributions across all three GO domains (cellular component, molecular function, and biological process) were assessed with the top 15 GO terms of each species visualized via ggplot2 in R. Shared and unique orthologous groups were visualized using UpSet plots, while protein domains and families were identified using InterProScan v5.75 [42].

### Comparative genomics

The *P. jefferyi* draft genome made by our study was compared to the previously submitted reference genome for the *P. jefferyi* in Genbank [19], as well as 21 other genomes of raptors under the family Accipitridae. Highly conserved orthologs in the BUSCO aves_odb10 database were extracted for each of the genome assemblies, and those common to all species were used to construct gene alignments using MAFFT v.7.5.0 [43, 44] and trimmed for terminal regions using trimAl v1.4.1 [45]. A total of 1,689 trimmed gene alignments were obtained from this and concatenated to generate an overall alignment. The alignment file was then filtered for polymorphic sites such as single-nucleotide polymorphisms (SNPs) and insertion-deletion mutations (indels) to construct the final phylip file that was used for phylogeny construction. Lastly, IQ-TREE2 v2.4.0 [46] was used to select for the best nucleotide substitution model and for the construction of the maximum-likelihood phylogeny with 1000 bootstraps. The reconstructed phylogenetic tree was visualized in iTOL [47].

### Heterozygosity and Demographic History

Individual reads were aligned to the meta-assembly using BWA-MEM2 v2.2.1, with resulting SAM files processed using SAMtools v1.16.1 [48, 49]. Alignment files were further refined using GATK v4.6.1 AddOrReplaceReadGroups and MarkDuplicates modules [50]. Folded site frequency spectrum was called in ANGSD v0.940 using -dosaf 1 and realSFS with the -minq 20 and -minmapq 30 filters [51]. Genome-wide heterozygosity for each individual was calculated as the ratio of heterozygous sites (H₁) to total sites from individual folded site frequency spectra [52].

Historical demographic patterns were inferred using Stairway Plot 2 with folded site frequency spectra [53]. Effective population size (Nₑ) changes were estimated assuming a mutation rate of 4.6 × 10⁻⁹ per site per generation (derived from collared flycatcher estimates; Smeds et al., 2016, and was used in vultures, Zou et al., 2021, and seagulls, Huynh et al., 2023) and a generation time of 18 years (IUCN Red List data) [54, 55, 56, 57]. Analysis employed the default two-epoch model with 67% sites sampled for training, 200 bootstrap replicates, and singleton exclusion. Historical Ne was estimated using the formula: θ = 4Neμ, where mean genome-wide heterozygosity served as a proxy for θ, with the same mutation rate, and mutation-drift equilibrium was assumed [58].

## Results

### Genome Assembly

Two meta-assemblies of the Philippine Eagle genome were created, one with Redbean and one with Flye. The total genome size was estimated to be 1.17 Gbp in Redbean and 1.16 Gbp in Flye. The Redbean and Flye meta-assemblies contained 6,149 and 10,732 contigs, respectively, greatly reducing the 156,269 average number of contigs from the 35 *P. jefferyi* de novo assemblies (Supplementary Table S2). The Redbean and Flye had N50 values of 395,632 and 217,643, respectively. Both Redbean and Flye assemblers were also able to eliminate the presence of ambiguous bases (Ns) in their meta-assemblies, with zero ‘#Ns per 100 kbp’ for both meta-assemblies. This is in contrast to the average de novo assemblies from the Philippine Eagle individuals, having 1,530 ambiguous bases per 100 kbp (Table 1). Overall, quality metrics of meta-assemblies from both methods showed great improvement of the Philippine Eagle genome compared to metrics of the initial de novo assemblies.

**Table 1.** Assembly metrics of average short-read assembly, draft meta-assembly, and NCBI reference genome [19] of *Pithecophaga jefferyi*.

| Assembly Metrics | Average Short-read Assembly | Meta-assembly | NCBI Reference genome |
| --- | --- | --- | --- |
| No. of contigs ($\geq 0$ bp) | 859,031 | 10,746 | 11,238 |
| No. of contigs ($\geq 1000$ bp) | 45,888 | 10,739 | 4,605 |
| No. of contigs ($\geq 5000$ bp) | 32,203 | 10,726 | 505 |
| Total length ($\geq 0$ bp) | 1,289,398,282 | 1,162,941,722 | 1,257,274,458 |
| Total length ($\geq 1000$ bp) | 1,174,997,506 | 1,162,937,479 | 1,252,843,518 |
| Total length ($\geq 5000$ bp) | 1,141,555,322 | 1,162,898,385 | 1,244,830,581 |
| No. of contigs | 54,511 | 10,732 | 1,135 |
| Largest contig | 461,899 | 1,384,840 | 84,365,990 |
| Total length | 1,181,066,427 | 1,162,923,170 | 1,247,235,770 |
| GC (%) | 41.82% | 41.56% | 41.37% |
| N50 | 55,606 | 217,643 | 38,333,288 |
| # N's per 100 kbp | 691.096 | 0 | 13,178.38 |
| BUSCOs | C:78.6% [S:78.3%,D:0.3%],<br>F:11.9%, M:9.5% | C:95.4%[S:94.9%,D:0.5%],<br>F:1.4%,M:3.2% | C:85.1%[S:84.8%,D:0.3%],<br>F:6.7%,M:8.2% |

Benchmarking Universal Single-Copy Orthologs (BUSCO) under the Aves_odb10 database showed higher completeness of the Flye meta-assembly (95.4%) compared to Redbean (90.1%). The Flye meta-assembly also had fewer missing BUSCOs (3.2%) than the Redbean one (9.1%). Notably, the redbean meta-assembly had comparable BUSCO completeness to the average de novo assemblies (90.1% vs 88.9%) and an increased portion of missing BUSCOs (9.1% vs. 4.8%) (Supplementary table S2). Because of this, the Flye meta-assembly was ultimately chosen as the working reference genome for downstream analysis in this study.

The available reference genome for *P. jefferyi* (GCA_025728025.1) was also compared to our draft meta-assembly. Based on the number of contigs, the largest contig, and the N50 of the reference genome, it appears to have a much greater contiguity than our draft genome meta-assembly. However, upon checking the proportion of unresolved gaps (13,178 #Ns per 100 kbp) and BUSCO completeness (85.1%) metrics of the NCBI reference genome, we see a great improvement in completeness in our draft genome, which is free of unresolved base calls and has a 95.4% BUSCO completeness. This also makes it more amenable for use in gene annotation in our downstream analysis [59, 60, 61].

Meanwhile, repeat masking was done before genome annotation and subsequent downstream analysis on heterozygosity and demographic history. RepeatMasker masked 59,132,847 bases (5.08%) of the genome, with interspersed repeats comprising the majority (Figure 1B). Long interspersed nuclear elements (2.41%), long terminal repeats (1.28%), and simple repeats (0.89%) accounted for the largest fractions of masked bases, alongside smaller contributions from DNA transposons (0.20%) and low complexity repeats (0.17%).

**Figure 1.**
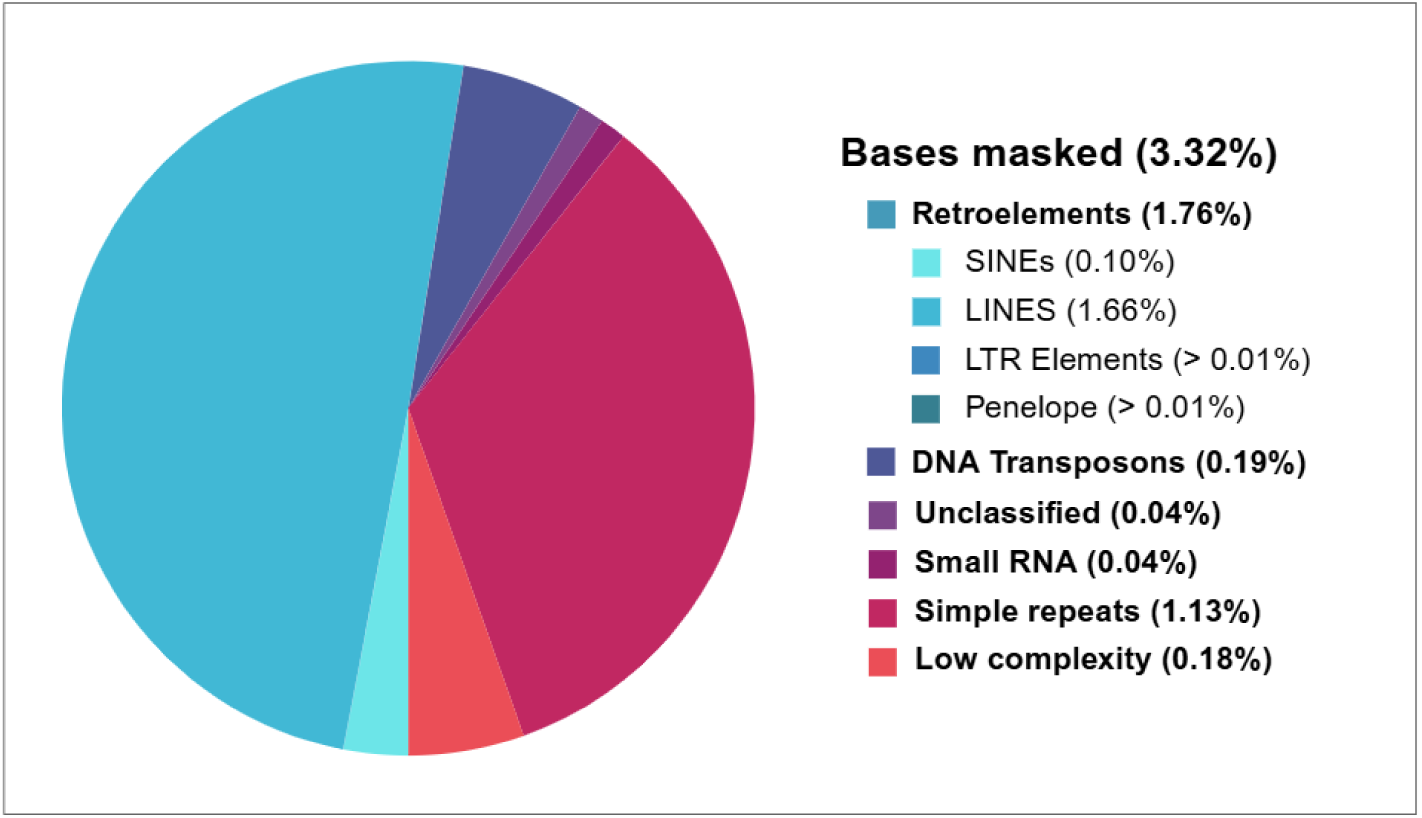
Repeat elements identified in the draft genome of *Pithecophaga jefferyi*.

### Genome Annotation

Before repeat masking, tRNA annotation performed using tRNAscan-SE identified a total of 191 tRNA genes confirmed by Infernal, comprising 170 canonical tRNAs, 17 predicted pseudogenes, 3 selenocysteine tRNAs, and 1 unclassified tRNA. Notably, 20 of the 64 possible anticodons were absent, including both ACA and GCA codons for cysteine, indicating a lack of corresponding tRNA genes for this amino acid. Ribosomal RNA annotation via barrnap revealed five rRNA features: partial sequences of 18S (27% alignment), 28S (31% alignment), and 5S (73% alignment), along with two complete 5S rRNA genes. No 5.8S rRNA genes were detected.

The final gene set produced by EVM comprises 24,781 genes (Table 2), spanning a total length of 347,818,533 base pairs and covering 29.90% of the draft assembly, with a gene density of approximately 21.30 genes per megabase. BUSCO analysis of the protein sequences revealed 93.0% completeness, with 2.8% fragmented and 4.2% missing genes, an improvement over the best pre-EVM annotation, GeMoMa, which showed 90.2% completeness, 1.4% fragmented, and 8.4% missing. In addition to increasing the number of complete BUSCOs, the final gene set significantly reduced the proportion of duplicate complete orthologous genes from 36.4% to just 0.5%, indicating a more refined and less redundant annotation. The higher duplication rate in GeMoMa is likely due to its use of homology-based predictions from three different Accipitridae raptor species.

**Table 2.** Structural annotation of protein-coding genes of *Pithecophaga jefferyi*.

|  | **EVM** | **GeMoMa** | **Aug-Raptor** | **Aug-Chicken** |
| --- | --- | --- | --- | --- |
| Total Gene Count | 24,781 | 30,404 | 27,822 | 46,461 |
| Total Gene Length (bp) | 347,818,533 | 772,824,705 | 366,529,379 | 426,324,956 |
| Mean Gene Length (bp) | 14,035 | 25,418 | 13,174 | 9,175 |
| Gene Density (genes/Mb) | 21.30 | 26.14 | 23.92 | 39.95 |
| Gene Coverage | 29.90% | 66.45% | 31.51% | 36.65% |
| Complete BUSCOs | 93.0% | 90.2% | 83.9% | 76.1% |
| Single-copy BUSCOs | 92.5% | 53.9% | 83.3% | 75.5% |
| Duplicated BUSCOs | 0.5% | 36.4% | 0.6% | 0.6% |
| Fragmented BUSCOs | 2.8% | 1.4% | 6.5% | 10.1% |
| Missing BUSCOs | 4.2% | 8.4% | 9.6% | 13.8% |

Out of the 24,781 genes identified, eggNOG-mapper functionally annotated 15,072 genes (60.8%), while InterProScan detected functional domains in 20,887 genes (84.3%). GO annotations at level 5, visualized in Figure 2A, highlighted the most frequently assigned terms across the three major GO categories: for Cellular Component, “intracellular membrane-bounded organelle” (65.4%), “nuclear lumen” (14.6%), and “cytoskeleton” (10.2%); for Molecular Function, “DNA binding” (14.1%), “kinase activity” (7.9%), and “regulatory region nucleic acid binding” (7.6%); and for Biological Processes, “cellular protein metabolic process” (21.5%), “regulation of gene expression” (20.1%), and “regulation of nucleobase-containing compound metabolic process” (18.9%); all reflecting evolutionarily conserved functions.

**Figure 2.**
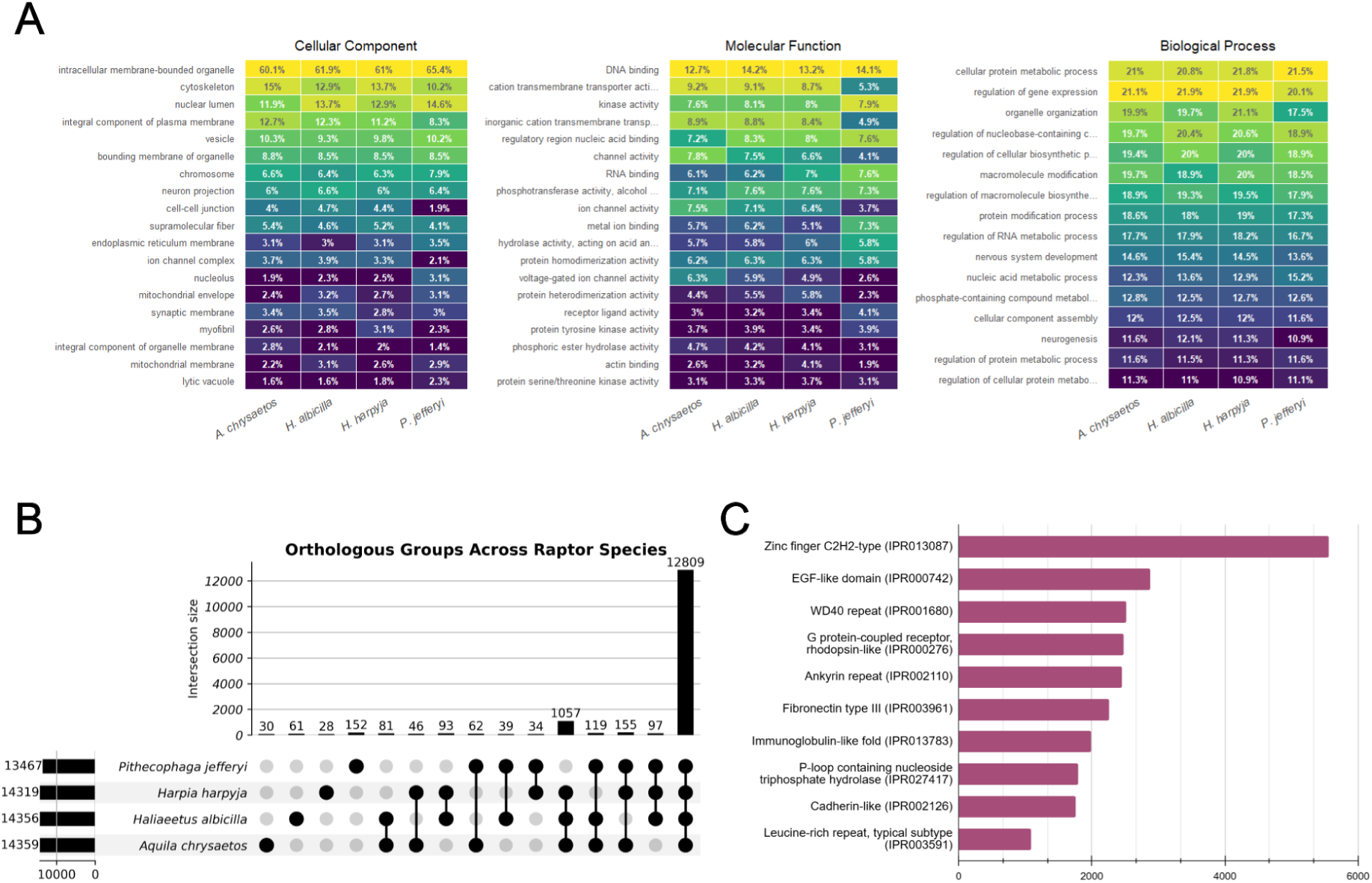
Summary of functional annotation of the Philippine Eagle compared to reference species. (A) Top GO term distributions at Level 5 across *A. chrysaetos, H. albicilla, H. harpyja,* and *P. jefferyi*, grouped under Cellular Component, Molecular Function, and Biological Process. (B) UpSet plot illustrating shared orthologous groups (OGs) among the four raptors. The bar graph counts group overlap, while the dot matrix indicates all possible species combinations. (C) Horizontal bar chart of the most represented InterProScan protein families.

**Figure 3.**
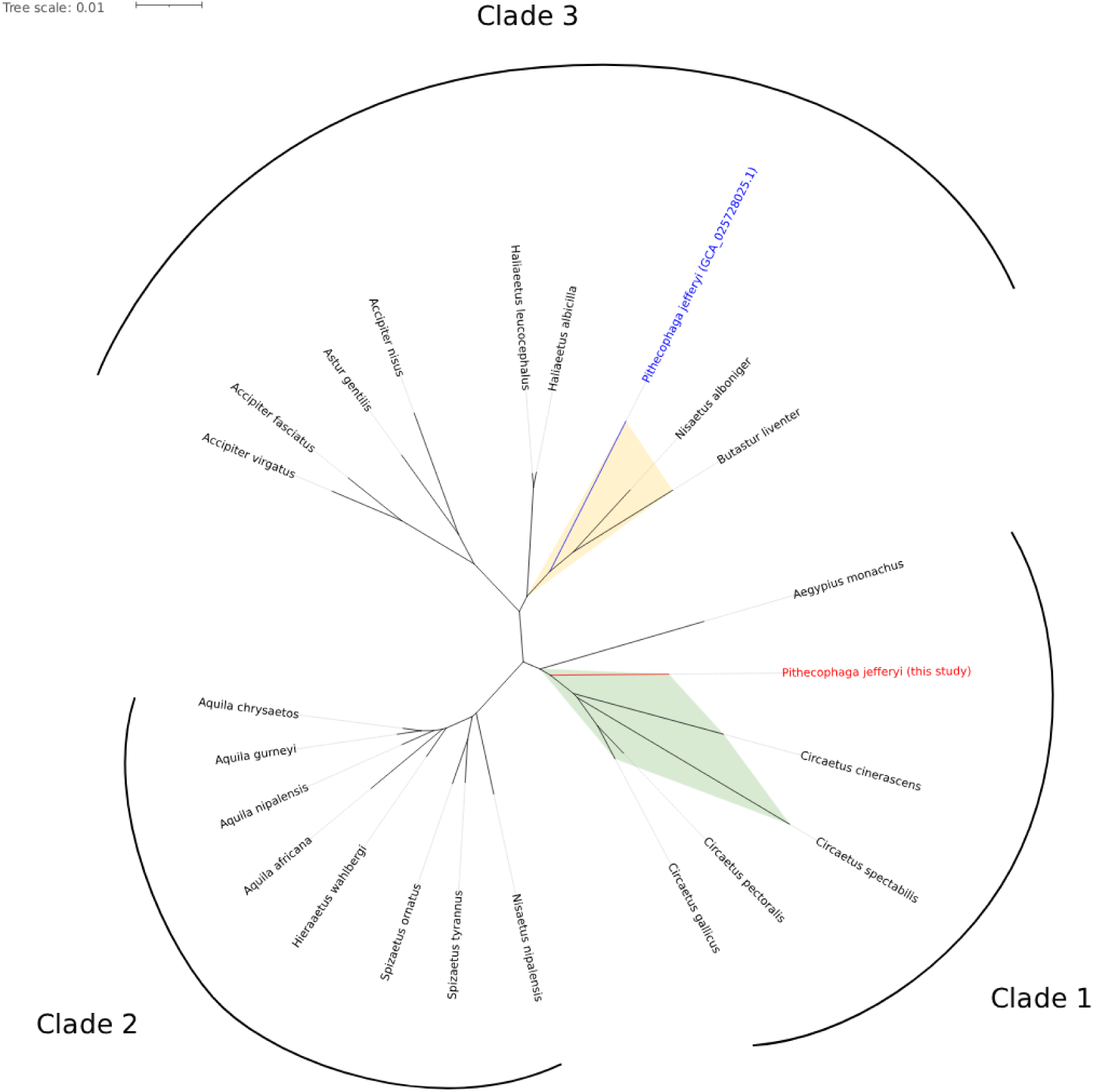
Phylogenetic tree of the draft genome of *P. jefferyi* made by this study, the *P. jefferyi* reference genome (GCA_025728025.1), and 21 raptor species under family Accipitridae. The phylip file used for phylogeny construction was derived from the polymorphic sites of concatenated alignments of universal single-copy orthologs. Major clades identified and labeled. Circaetinae species are shaded in light green, including the *P. jefferyi* draft genome from this study, labeled as “*Pithecophaga jefferyi* (this study),” and are highlighted in red text. The reference genome in NCBI is denoted as "*Pithecophaga jefferyi* (GCA_025728025.1)," is labeled with blue text and shaded in yellow along with *B. liventer* (GCA_026109245.1), and *N. alboniger* (GCA_025447895.1) to emphasize their misplacement away from their recognized subfamily in the Accipitridae.

These top terms were consistent with those found in other raptor proteomes. However, the Philippine Eagle showed notably lower representation of “cation transmembrane transporter activity,” “inorganic cation transmembrane transporter activity,” and related ion channel functions. EggNOG-mapper also assigned 13,467 orthologous groups (OGs) to *P. jefferyi*, with approximately 14,300 OGs in the reference proteomes; of these, 12,809 OGs (about 95%) were shared, while 1,057 OGs present in the reference proteomes were absent in *P. jefferyi*, indicating some missing core raptor annotations (Figure 2B).

Interestingly, *P. jefferyi* had 152 unique OGs, more than the 30, 61, and 28 unique OGs found in the other raptors. InterProScan further identified protein families and domains from various databases, shown in Figure 2C, with the most abundant being “Zinc finger C2H2-type” (IPR013087), “EGF-like domain” (IPR000742), and “WD40 repeat” (IPR001680), all of which are evolutionarily conserved.

### Comparative Genomics

The phylogenetic tree composed of the *P. jefferyi* draft genome, the currently available *P. jefferyi* reference genome (GCA_025728025.1), and 21 other raptors of the family Accipitridae (Figure 1, Supplementary table S3) showed 3 major clade formations. Phylogenetic placement of the draft genome in this study is within the clade of represented raptors from Circaetinae, which mirrors the taxonomic revisions made by the International Ornithological Congress, which also places *P. jefferyi* together with Snake eagles forming subfamily Circaetinae (Accipitridae, Accipitriformes) [63]. The *Aegypius monachus* genome also formed a clade with the subfamily Circaetinae. The genomes of *Nisaetus nipalensis*, *Spizaetus ornatus* and *tyrannus*, *Hieraeetus wahlbergi*, and the *Aquila* genus formed another clade, with species from *Aquila* exhibiting monophyly. The *Accipiter*, *Astur*, and *Haliaeetus* genera also formed a clade, and interestingly, *Nisaetus alboniger*, *Butastur liventer*, and the *P. jefferyi* reference genome (GCA_025728025.1) were also placed under this major clade.

### Genetic Diversity

The average genome-wide heterozygosity across all sampled Philippine Eagles is estimated at 0.000309. When compared across conservation statuses, heterozygosity values were consistently lower in species of higher threat categories, with Near Threatened, Vulnerable, Endangered, and Critically Endangered species showing significantly reduced levels relative to Least Concern taxa (Figure 4A). Among these, the Philippine Eagle exhibited one of the lowest heterozygosity values compared to the other Critically Endangered species.

**Figure 4.**
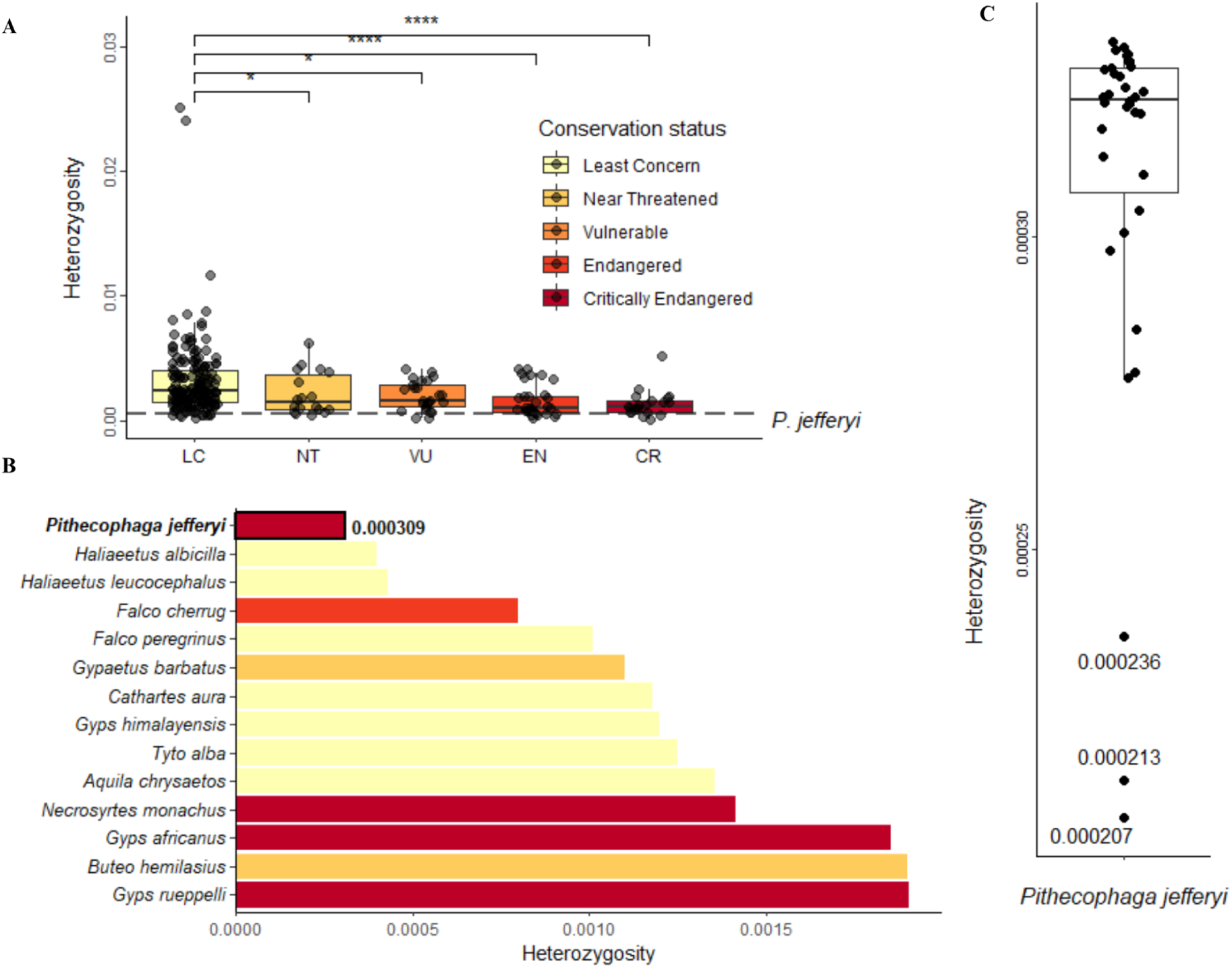
Genome-wide heterozygosity of the Philippine Eagle relative to other threatened species. (A) Boxplot distribution of genome-wide heterozygosity coefficients for threatened species, grouped by conservation status [52, 62]. The heterozygosity of the Philippine Eagle is demarcated with a dashed line. (B) Genome-wide heterozygosity of the Philippine Eagle relative to other threatened raptors (Accipitriformes, Strigiformes, Falconiformes, and Cariamiformes) [55]. (C) individually-calculated genome-wide heterozygosity for each eagle sample.

Focusing on raptors (from families belonging to avian orders Accipitriformes, Strigiformes, Falconiformes, Cathartiformes, and Cariamiformes), heterozygosity values were generally low across all conservation statuses, including those species listed as Least Concern [64]. The Philippine Eagle has one of the lowest genome-wide heterozygosity compared to raptors with available heterozygosity estimates.

### Demographic history

The inferred historic effective population size from the average genome-wide heterozygosity was 16,795, drastically larger than any population estimates today [4]. The Stairway plot for the Philippine Eagle, assuming the same mutation rate and a generation time of 18 years, suggested two major historical bottlenecks followed by an ongoing population decline (Figure 5). The earliest bottleneck occurred approximately 800,000 years ago, reducing the effective population size from ∼40,000 to 10,000 individuals. The population subsequently recovered to ∼100,000 individuals 400,000 years later. A second, more severe bottleneck began around 80,000 years ago, reducing Ne to its second lowest point of ∼4,000 individuals at approximately 50,000 years ago, coinciding temporally with the Youngest Toba Eruption (60,000 −80,000 years ago) as highlighted in the plot [65]. Following this bottleneck, the population recovered to ∼70,000 individuals by 30,000 years ago, corresponding to the Last Glacial Maximum, presumably when Mindanao, Samar, Leyte, and nearby islands formed a single landmass [66, 67].

**Figure 5.**
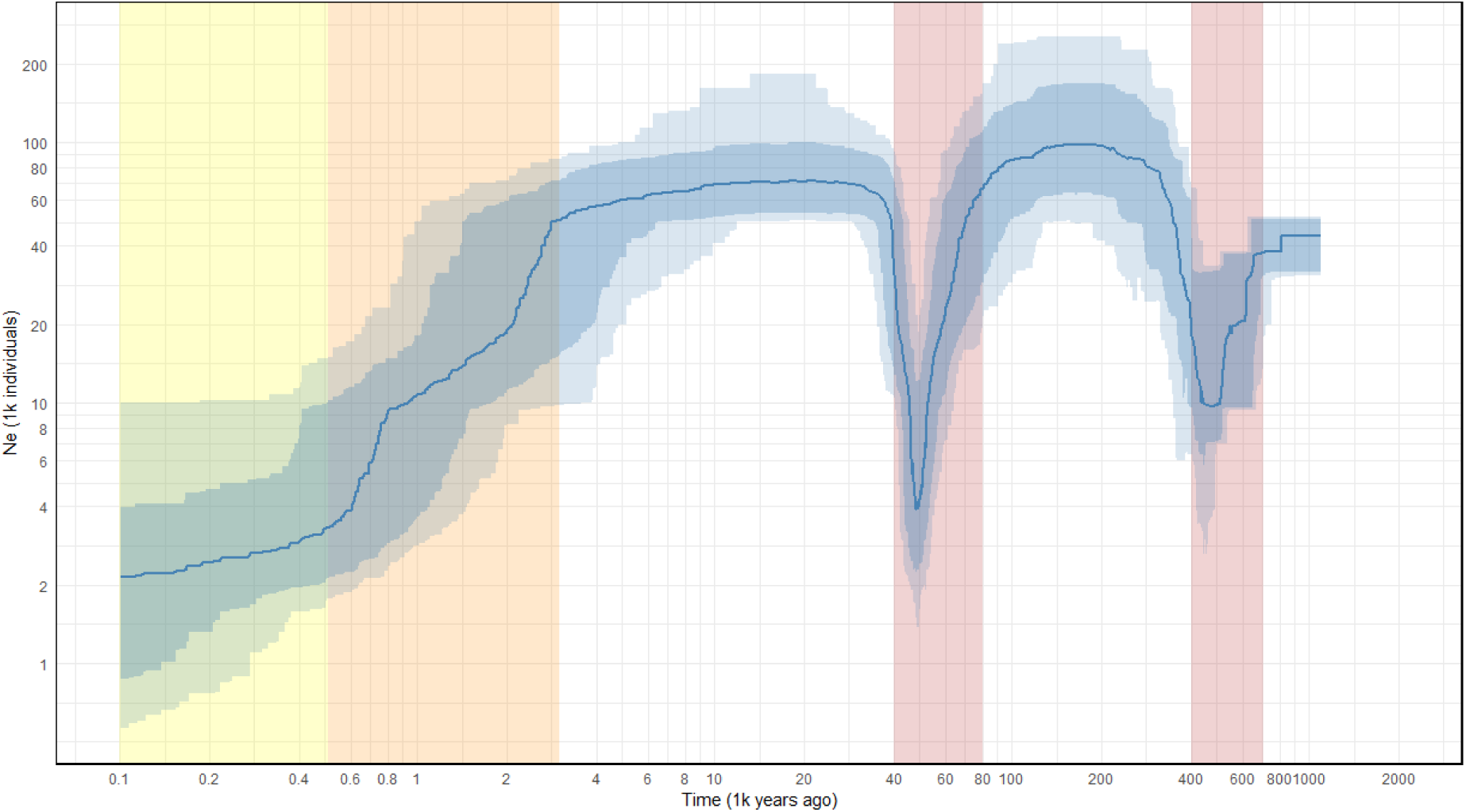
Stairway plot of the demographic history of the Philippine Eagle. The dark blue line represents the median estimate of effective population size over time. Blue and light blue shaded areas indicate the 87.5%–12.5% and 97.5%–2.5% confidence intervals, respectively. The red boxes highlight two population bottlenecks, the orange box highlights the start of the ongoing decline, and the yellow box highlights the start of deforestation in the Philippines.

The most concerning trend is an ongoing population decline that began approximately 3,000 years ago, with Ne decreasing from ∼50,000 to an estimated ∼2,000 individuals 100 years ago. This decline encompasses the period of Spanish colonization (500 years ago), when the Philippines maintained approximately 90% forest cover, contrasting sharply with the current 23% (72,000 km^2^) forest coverage [68, 69]. However, the population decline predates this significant deforestation by approximately 2,500 years, suggesting that forest cover loss alone cannot fully explain the Philippine Eagle’s current demographic trajectory. The ongoing decline also appears to coincide with the Austronesian expansion into the Philippine archipelago (∼2200-1000 BCE), suggesting a possible additional explanation [70, 71].

## Discussion

### Quality and Completeness of the Philippine Eagle Draft Assembly

The genome assembly for *P. jefferyi* generated in this study demonstrates improved quality compared to previous efforts, particularly when considering assembly methods, the impact of fragmented DNA, and the benefits of combining multiple assemblies into a consensus meta-assembly.

The meta-assemblies show significant enhancements in contiguity and completeness compared to individual assemblies. This approach increased genome coverage and effectively resolved ambiguous base calls. The meta-assemblies from both Redbean and Flye notably eliminated the presence of ambiguous bases (Ns). This signifies that unresolved regions in one individual assembly were complemented by contigs from other assemblies where those regions were resolved [72, 73, 74]. The final meta-assemblies also improved contiguity, indicated by higher N50 values.

The draft genome of *P. jefferyi* produced in this study demonstrates higher completeness and lower ambiguity when directly compared to the reference genome previously published in NCBI. The NCBI reference assembly was found to have a high proportion of unresolved gaps (N’s per 100 kbp), despite having a higher overall N50 value than the meta-assembly. The assembly was reportedly based on DNA extracted from a toe pad sample, which is generally recognized as being more degraded than DNA obtained from fresh tissue [75, 76]. This might have caused the high proportion of ambiguous base calls in the reference assembly, resulting in a lower BUSCO completeness score.

Conversely, the present study utilized DNA extracted from blood samples of live Philippine Eagles, providing higher-quality input material with detailed documentation. The meta-assembly pipeline used in the present study then addressed the inherent limitations of short-read de novo assemblies. Consequently, apart from the elimination of ambiguous base calls, the *P. jefferyi* draft genome from this study boasts a BUSCO completeness of 95.4%. These strong metrics represent a significant improvement for critical downstream applications like gene annotation and diversity analysis.

While the draft genome lays essential groundwork, its limitations highlight clear opportunities for future refinement through long reads and Hi-C sequencing to achieve chromosome-level resolution and transcriptomic validation to refine gene models and functional annotations. The difficulty in resolving repetitive regions in short read assemblies has led to incomplete annotation of rRNA genes, particularly 18S, 28S, and the missing 5.8S, due to their location in highly repetitive nucleolar organizer regions, while some of the shorter and more dispersed 5S rRNA were recovered [77]. The genome shows only 5.08% masked repeat sequences, significantly lower than the 16.33% in the Harpy Eagle, *Harpia harpyja*, likely due to collapsed repetitive regions inherent to short-read sequencing [78].

Functional annotation reveals a deficit of 1,057 orthologous groups (OGs) compared to reference raptors, suggesting an underestimation of the true gene set, though conserved biological pathways appear adequately represented. The 152 unique OGs may reflect either lineage-specific traits or false predictions from assembly fragmentation, which warrants further investigation [79]. While homology-based and ab initio methods provide a comparative framework, transcriptomic data will be essential to validate gene function, identify pseudogenes, and uncover splicing diversity [80, 81, 82]. Despite these limitations, this draft genome marks a foundational step for conservation genomics in Philippine raptors, enabling studies on population structure, adaptive evolution, and genetic monitoring, critical for managing the country’s rich biodiversity [83, 84, 85].

### Comparative Genomics in the Accipitridae family

The study found that the *P. jefferyi* draft genome formed a clade with raptors in the subfamily Circaetinae and *Aegypius monachus* (Cinereous Vulture, an Old World vulture) [86]. The placement of the draft genome with Circaetinae is consistent with the taxonomic revisions set by the International Ornithological Congress [63] and several prior molecular phylogenetic studies [87, 88]. Some more recent, larger-scale phylogenomic studies [89], using ultraconserved elements (UCEs), also placed *P. jefferyi* as sister to the Circaetus + Terathopius clade within Circaetinae.

The Circaetinae subfamily includes serpent eagles (*Spilornis*, 6 species), Bateleur (*Terathopius ecaudatus*), Snake Eagles (Circaetus, 7 species), and Madagascan Serpent Eagle (*Eutriorchis ater*) [63]. This relationship of the draft genome (this study) with Circaetinae supports the taxonomic origins of the extremely large *Pithecophaga* to be derived from smaller ancestral snake eagles. Hence, supporting the result of studies on the diet of *P. jefferyi,* which comprises 25% snakes and up to 37% reptiles [90]. This taxonomic arrangement also confirms that *P. jefferyi* is not related to harpy eagles belonging to the subfamily Harpiinae, which includes the enormous Harpy Eagle (*Harpia harpyja*) and Papuan Harpy Eagle (*Harpyopsis novaeguineae*).

The grouping of Old World vultures (Aegypiinae/Gypaetinae) and snake eagles (Circaetinae) has also been a consistent finding in many analyses [85, 86], often showing them as sister groups or within a larger clade. Additionally, the phylogeny correctly resolved the booted eagles (Aquilinae), which include the *Nisaetus*, *Spizaetus*, *Hieraaetus*, and *Aquila* genera [91, 87, 88]; sea eagles (Haliaeetinae), which consist of the *H. leucocephalus* and *H. albicilla* eagles [87, 93], and goshawks (Accipitrinae), consisting of the *Accipiter* and *Astur* raptors [89, 93]. The presence of these empirically consistent phylogenetic placements further reinforces the robustness of the BUSCO-based analysis.

There were also phylogenetic inferences based on the reference genomes for *P. jefferyi* (GCA_025728025.1), *B. liventer* (GCA_026109245.1), and *N. alboniger* (GCA_025447895.1) that resulted in an alternative, known conflicting topology, placing it within the *Accipiter*, *Astur*, *Haliaeetus* clade. This observed topological discordance may reflect challenges in genome assembly and scaffolding when utilizing short-read sequences derived from highly fragmented DNA, particularly in repeat-rich regions [89, 94, 95], and imply that the quality of these available genome assemblies warrants further assessment. The robust placement of *P. jefferyi* into the Circaetinae clade achieved by our draft genome successfully resolves this phylogenetic inconsistency, consistent with established molecular evidence [63, 87, 88, 89]. Despite a simple BUSCO-based comparative phylogenetic methodology, the empirically consistent placement of the *P. jefferyi* draft genome indicates that the assembly may be a more accurate representation to the biological reality of the Philippine Eagle genome compared to the available NCBI reference.

### Demographic History Reveals Ancient Bottlenecks and a Recent Ongoing Decline

The effective population size (Ne) provides insight into the demographic history and genetic health of a species, representing the number of individuals in an idealized population that would experience the same rate of genetic drift as the observed population [96]. In this study, the Ne was inferred using the site frequency spectrum, which estimates a long-term historical average over many generations. Accordingly, the resulting Ne represents the size of a hypothetical stable population maintained throughout the species’ existence, rather than its current demographic status.

For the Philippine Eagle, the inferred historic effective population size in this study was approximately 17,000, which is larger than current census population estimates of around 392 breeding pairs [4]. This large difference between the effective and census (Nc) population size suggests that the population is undergoing a relatively recent and steep decline. While Ne is generally expected to be lower than Nc in stable populations, owing to factors such as unequal reproductive success and non-random mating, the reversal observed here suggests a demographic contraction from a historically larger, more genetically diverse population [97]. In effect, the elevated Ne serves as genetic evidence for a trend already recognized through field observations – the Philippine Eagle population is shrinking and remains at significant risk.

Similar patterns have been observed in other raptor species, such as Golden Eagle (*Aquila chrysaetos*) subpopulations in Japan and Scotland, where historical Ne estimates of ∼70,000 likely predate population isolation and reflect the effective size of larger, geographically proximate continental populations before fragmentation [98]. For the Philippine Eagle, which is found across multiple islands (Luzon, Leyte, Samar, and Mindanao), the high historical Ne estimate likely corresponds to a time before island fragmentation and subsequent population decline. It should be noted that the samples analyzed in this study were predominantly from Mindanao, with one individual from Samar; therefore, they may not fully capture the genetic diversity across all subpopulations, particularly those from Luzon and Samar. The Ne estimate likely reflects a long-term average that predates the current fragmented island distribution and recent population contractions.

The reconstructed demographic history for the Philippine Eagle reveals two significant bottlenecks and an ongoing population decline. The most recent bottleneck began around 80,000 years ago, reducing the effective population size (Ne) to approximately 4,000 individuals. This decline coincided with the Youngest Toba Eruption that occurred around 74,000 years ago [65]. The eruption happened at what is now Lake Toba in Sumatra, Indonesia, which lies in roughly the same geographical region as the present distribution of the Philippine Eagle [65]. While the spatial and temporal coincidence of this volcanic eruption with the inferred bottleneck suggests a possible association, it does not provide definitive evidence of causation. The Youngest Toba Eruption was initially hypothesized to have caused severe climatic impacts and population bottlenecks across various species, including humans [99]. However, more recent studies suggest that the eruption’s effects on global climate may have been overestimated [65]. This raises the possibility that factors beyond the Toba eruption may have contributed to the Philippine Eagle’s demographic decline during this period.

The most recent bottleneck fits within broader avian demographic trends during the Last Glacial Period (LGP; 110,000 years ago). Demographic reconstructions using whole genomes and PSMC analysis revealed that many bird species experienced significant effective population size reductions during this period, with 22 of 38 analyzed species showing notable declines [100]. Raptors like the White-tailed Eagle (*Haliaeetus albicilla*), Bald Eagle (*Haliaeetus leucocephalus*), Barn Owl (*Tyto alba*), and Turkey Vulture (*Cathartes aura*) exhibited sustained population collapses, with the eagles declining from pre-LGP levels of ∼60,000 individuals to post-glacial lows of only 2,000-6,000 individuals by 10,000 years ago [100, 101].

In contrast, however, the Philippine Eagle’s effective population size (Ne) rebounded to approximately 70,000 individuals as early as 30,000 years ago, approaching pre-LGP estimates and exhibiting a recovery trajectory not observed in other raptors [100]. This demographic resurgence may be partially attributed to the formation of the Greater Mindanao faunal region or the Mindanao Pleistocene Aggregate Island Complex (PAIC) during the LGP, when sea levels fell by approximately 120-130 meters. This eustatic drop exposed extensive continental shelves and land bridges that temporarily unified Mindanao, Samar, Leyte, Bohol, Dinagat, and adjacent satellite islands into a single contiguous landmass, as delineated by Heaney’s faunal regions and the PAIC model using the 120-meter bathymetric line [66, 67, 102]. These resulting land aggregates substantially expanded the total land area of the Philippine archipelago — potentially reaching ∼350,000–400,000 km² — thereby increasing habitat availability, potentially enhancing prey abundance, and facilitating gene flow among subpopulations that had been isolated during previous interglacial periods.

Notably, even during the Last Glacial Maximum (∼22,000 years ago), Ne estimates indicate that the Philippine Eagle population persisted, in contrast to the sustained declines observed in other avian populations, suggesting a degree of resilience to climatic challenges, potentially mediated by the ecological buffering effects of PAIC-driven habitat aggregation and landmass expansion.

Although the species demonstrated resilience during earlier climatic fluctuations, its demographic history over the past 3,000 years reflects a prolonged and ongoing population decline, suggesting that the hypothesized ecological advantages conferred by the Greater Mindanao faunal region or Greater Mindanao PAIC had since been lost, as Holocene sea levels disaggregated these landmasses by ∼7,000 years ago [103]. This reduction in landmass and loss of habitat connectivity, compounded by emerging stressors such as early anthropogenic pressures, may have disrupted population stability. Effective population size estimates indicate a drop from ∼50,000 to ∼2,000 individuals between 3 kya and 0.1 kya, which is faster and more severe than earlier bottlenecks. While deforestation is a recognized anthropogenic threat, it can only partially explain the timing of the population decline. Spanish colonization of the Philippines began 500 years ago, when forest cover was still at 90%, declining to 71% by 100 years ago (around 1900), coinciding with the endpoint of our Ne estimates [14, 68]. Forest cover has since been drastically reduced to only 23% (72,000 km^2^) by 2002, of which less than 10,000 km^2^ are considered primary forests [69, 104]. This timeline reveals that a significant population decline was already well underway even before major deforestation began, suggesting the influence of additional stressors, may it be ecological, climatic, or other anthropogenic causes. Notably, the onset of this decline also coincides with the Austronesian expansion into the Philippine archipelago (∼2200–1000 BCE), a period characterized by increasing human population densities, intensified hunting activities, and heightened competition for key eagle prey species such as wild pigs, deer, and macaques. These ecological pressures may have contributed significantly to the species’ long-term population contraction, suggesting another potential anthropogenic influence on the population history of the Philippine Eagle. [70, 71].

This dissonance between known anthropogenic threats and the timing of population decline stands in stark contrast to the demographic trajectory of other critically endangered birds, such as the Crested Ibis (*Nipponia nippon*), whose recent population collapse due to overhunting and habitat loss was clearly captured in demographic reconstructions [62]. Unlike the Philippine Eagle, whose decline spans millennia, the Crested Ibis experienced a rapid reduction from ∼10,000 individuals to near extinction within the last 100 years, closely aligned with documented anthropogenic pressures. Although our Stairway Plot cannot resolve Ne trends beyond ∼100 years ago, and therefore cannot determine whether populations have stabilized since conservation efforts began in 1969, the Philippine Eagle’s long-term decline remains clearly detectable [11]. Recent threats such as habitat loss and human persecution are well documented, yet the prolonged nature of the Philippine Eagle’s decline raises the possibility that older, less understood ecological or anthropogenic pressures may have also played a role.

### Extremely Low Genetic Diversity and Drift Debt Threaten Long-term Viability

The genomic consequences of the ongoing long-term demographic decline are evident in the Philippine Eagle’s exceptionally low genome-wide heterozygosity (GWH). Among all raptors with reported GWH estimates, the Philippine Eagle exhibits the lowest value, at just 0.000309. Although low heterozygosity alone does not determine extinction risk, since some species may persist with low diversity and others may be endangered despite moderate heterozygosity (see Figure 4A), this value is concerning when interpreted alongside the species’ sustained small population size. With an estimated 392 breeding pairs remaining in the wild, and some more pessimistic assessments suggesting as few as 64 individuals, prolonged genetic drift and limited opportunities for gene flow have likely contributed to the erosion of genetic variation [4, 105]. In this context, its low heterozygosity aligns with well-documented demographic decline and ecological constraints. As such, it serves as a genomic signal of heightened risk, with important implications for the species’ persistence and adaptive potential.

In small populations, reduced genome-wide heterozygosity indicates a history of strong genetic drift, which reduces the efficacy of purifying selection against mildly deleterious alleles [106]. Theoretical expectations also suggest that limited mate choice in such populations increases the likelihood of inbreeding, which can elevate homozygosity and expose recessive harmful variants [107]. Although inbreeding was not directly assessed in this study, these dynamics are consistent with the observed low heterozygosity and raise concerns about potential inbreeding depression, reduced fitness, and long-term viability [106, 108]. Genomic resources presented in this study may inform future breeding and management programs, with the protection of the remaining genetic variation being critical to their success. Conservation efforts must extend beyond short-term demographic recovery to ensure both genetic resilience and long-term persistence.

While the Philippine Eagle requires short-term intensive population recovery efforts, its critically low heterozygosity raises questions about whether demographic gains alone will ensure long-term persistence. As the largest island-endemic raptor, a combination of traits predicting elevated conservation status, the species faces compounded challenges in achieving genetic resilience [109]. Comparative examples from both large continental raptors and island-endemic species illustrate this pattern of genetic vulnerability despite demographic recovery. Large continental eagles such as the White-tailed Eagle (*Haliaeetus albicilla*) and the Bald Eagle (*Haliaeetus leucocephalus*) were downlisted to Least Concern following population recovery from DDT-driven declines, yet both retain low genome-wide heterozygosity [110, 111, 112, 113]. Among island-endemic raptors, the endangered Mauritius Kestrel (*Falco punctatus*) recovered from a near-extinction event of four remaining individuals to now around 140 individuals; despite demographic recovery, genomic analysis from a recent preprint revealed it to have critically low heterozygosity and evidence of sustained recent inbreeding [114, 115, 116]. More broadly, island-endemic birds show similar patterns: the Chatham Island Robin (*Petroica traversi*) was downlisted from Endangered to Vulnerable following recovery from a single breeding pair, and consequent genomic analysis revealed substantial decreases in heterozygosity and increases in F_ROH_ from historical samples [117, 118, 119]. A similar trajectory is observed in the Seychelles Magpie-Robin (*Copsychus sechellarum*), an island-endemic passerine that recovered from a severe bottleneck of just 12 individuals in the 1960s through translocation-based conservation efforts and was downlisted from Critically Endangered to Endangered in 2005, despite the absence of comprehensive genetic data at the time [120]. Subsequent genomic analysis in 2021 measured its genome-wide heterozygosity at 0.000150, around half of that of the Philippine Eagle, and observed long runs of homozygosity, indicating recent inbreeding [121]. More troublingly, they noted that funding and research efforts plateaued following the downlisting, demonstrating how premature declarations of conservation success can compromise long-term protection when genetic factors are overlooked.

These examples demonstrate that demographic recovery does not erase genomic consequences of bottlenecks. The Philippine Eagle faces the additional challenge of ‘drift debt’ – even with population size recovery, the species would remain genetically imperilled due to genomic erosion from past bottlenecks, with genetic drift effects persisting for many generations [99, 122]. For the Philippine Eagle, genetic considerations must be integrated in any future conservation assessment to prevent similar outcomes to the Seychelles Magpie-Robin, where downlisting preceded comprehensive genetic analysis and led to reduced conservation investment, leaving the species genetically exposed to future threats.

## Conclusion

We have successfully constructed and annotated a draft genome for the Philippine Eagle that improves on the completeness and genome coverage of the previously established reference genome. Comparative phylogenetic analysis then confirmed its placement within the subfamily Circaetinae, something the previous reference genome was not able to reflect. This clade placement with snake eagles also agrees with the taxonomic revisions of the International Ornithological Congress.

By using this genomic resource, we found that the Philippine Eagle has one of the lowest genome-wide heterozygosities, at an average of 0.000309, compared to raptors and other threatened species. We also found that the Philippine Eagle has gone through two population bottlenecks and an ongoing population decline that predates deforestation in the Philippines. Its critically low heterozygosity limits its adaptive capacity and, combined with its dwindling population, makes the Philippine Eagle one of the most vulnerable species to ongoing anthropogenic and environmental pressures. Conservation efforts could utilize this genomic resource in monitoring the success of captive breeding and translocation programs.

## Authors’ contributions

Conceptualization: DMP and CPS; sample collection: JI; methodology and formal analysis: KP, SMA, JL, MCA, RJM, JME, CL, DMP, and FP; writing—original draft: KP, SMA, JL, MCA, DMP, FP, CL, and FT; writing—review and editing, all authors; supervision: FT, JCG, JI, and CPS.

## Acknowledgements

We would like to thank the biologists and volunteer veterinarians at the Philippine Eagle Foundation Inc. for providing Philippine Eagle samples, the University of the Philippines System -Philippine Genome Center for partial funding support, and MGI for providing DNBSEQ sequencing kits. The PEF has a Memorandum of Agreement with the Department of Environment and Natural Resources (DENR) for research and conservation of Philippine Eagles across the country.

