## Supplementary Tables 1-3 for "Genomic Analysis Reveals Recent Population Decline and Exceptionally Low Genome-Wide Heterozygosity of the Critically Endangered Philippine Eagle, *Pithecophaga jefferyi* (Aves: Accipitridae)"

### Supplementary Materials

**Table S1.** Details of the sampled Philippine Eagles

| Sample | Day hatched or rescued | Province |
|---|---|---|
| PEF 01 | 11/23/2006 | Davao del Sur |
| PEF 02 | 9/25/2021 | Davao de Oro |
| PEF 03 | 6/28/2019 | Eastern Samar |
| PEF 04 | 8/28/2020 | Davao Oriental |
| PEF 05 | 4/13/2007 | Surigao del Sur |
| PEF 06 | 2/12/2022 | Lanao del Sur |
| PEF 07 | 2/11/2000 | Davao del Sur |
| PEF 08 | 12/18/2004 | Davao del Sur |
| PEF 09 | 4/25/2015 | Lanao del Sur |
| PEF 10 | 2/23/1999 | Davao del Sur |
| PEF 11 | 2/9/2013 | Davao del Sur |
| PEF 12 | 5/7/2003 | Sarangani |
| PEF 13 | 11/25/2005 | Davao del Sur |
| PEF 14 | 2/26/2021 | Bukidnon |
| PEF 15 | 3/7/2010 | Davao del Sur |
| PEF 16 | 1/12/2011 | Davao Oriental |
| PEF 17 | 1/16/2003 | Davao del Sur |
| PEF 18 | 11/7/2015 | Davao del Sur |
| PEF 19 | 2/4/2002 | Davao del Sur |
| PEF 20 | 11/11/1993 | Sarangani |
| PEF 21 | 2/24/2023 | Bukidnon |
| PEF 22 | 5/5/1984 | Misamis Oriental |
| PEF 23 | 11/11/2000 | Davao del Sur |
| PEF 24 | 4/18/2011 | Davao del Sur |
| PEF 25 | 1/2/2024 | Davao del Sur |
| PEF 26 | 10/25/1992 | Davao del Sur |
| PEF 27 | 12/7/2007 | Davao del Sur |
| PEF 28 | 12/15/2000 | Davao del Sur |
| PEF 29 | 9/15/2002 | Agusan Del Norte |
| PEF 30 | 3/31/2022 | Agusan del Sur |
| PEF 31 | 8/11/2023 | Davao del Sur |
| PEF 32 | 11/22/2002 | Zamboanga del Norte |
| PEF 33 | 12/10/2001 | Davao del Sur |

| | | |
|---|---|---|
| PEF 34 | 2/24/2024 | Bukidnon |
| PEF 35 | 7/1/1997 | Misamis Occidental |

**Table S2.** Assembly metrics of Flye and Redbean meta-assemblies for *Pithecophaga jefferyi*

| Assembly Metrics | Flye Meta-assembly | Redbean Meta-assembly |
|---|---|---|
| No. of contigs (>= 0 bp) | 10,746 | 6,149 |
| No. of contigs (>= 1000 bp) | 10,739 | 6,149 |
| No. of contigs (>= 5000 bp) | 10,726 | 6,139 |
| Total length (>= 0 bp) | 1,162,941,722 | 1,168,224,302 |
| Total length (>= 1000 bp) | 1,162,937,479 | 1,168,224,302 |
| Total length (>= 5000 bp) | 1,162,898,385 | 1,168,176,638 |
| No. of contigs | 10,732 | 6,149 |
| Largest contig | 1,384,840 | 2,550,894 |
| Total length | 1,162,923,170 | 1,168,224,302 |
| GC (%) | 41.56 | 41.11 |
| N50 | 217,643 | 395,632 |
| # N's per 100 kbp | 0 | 0 |
| BUSCOs | C:95.4%[S:94.9%,D:0.5%],F:1.4%,M:3.2% | C:90.1%[S:88.1%,D:2.0%],F:0.8%,M:9.1% |

**Table S3.** Pertinent information on raptor genome assemblies used for comparative genomics analysis.

| Species | Genome Accession | Isolation source | Sequencing method | Submitter |
|---|---|---|---|---|
| *Accipiter fasciatus* | GCA_027475485.1 | muscle | Illumina HiSeqX | Iridian Genomes |
| *Accipiter nisus* | GCA_027497715.1 | muscle | Illumina HiSeqX | Iridian Genomes |
| *Accipiter virgatus* | GCA_025728005.1 | toe pad | Illumina HiSeqX | Iridian Genomes |
| *Aegypius monachus* | GCA_025309335.1 | toe pad | Illumina HiSeqX | Iridian Genomes |
| *Aquila africana* | GCA_027497815.1 | toe pad | Illumina HiSeqX | Iridian Genomes |
| *Aquila chrysaetos* | GCF_900496995.4 | heart muscle tissue | PacBio, Illumina, BioNano, Dovetail | Wellcome Sanger Institute |
| *Aquila gurneyi* | GCA_034782695.1 | tissue | Illumina HiSeqX | Iridian Genomes |

| | | | | |
|---|---|---|---|---|
| *Aquila nipalensis* | GCA_024363155.1 | toe pad | Illumina HiSeqX | Iridian Genomes |
| *Astur gentilis* | GCF_929443795.1 | heart tissue | PacBio, Illumina, Arima2 | Wellcome Sanger Institute |
| *Butastur liventer* | GCA_026109245.1 | toe pad | Illumina HiSeqX | Iridian Genomes |
| *Circaetus cinerascens* | GCA_026770385.1 | toe pad | Illumina HiSeqX | Iridian Genomes |
| *Circaetus gallicus* | GCA_025504725.1 | tissue | Illumina HiSeqX | Iridian Genomes |
| *Circaetus pectoralis* | GCA_035590135.1 | tissue | Illumina HiSeqX | Iridian Genomes |
| *Circaetus spectabilis* | GCA_034781195.1 | toe pad | Illumina HiSeqX | Iridian Genomes |
| *Haliaeetus albicilla* | GCF_947461875.1 | blood sample | PacBio, Arima2 | Wellcome Sanger Institute |
| *Haliaeetus leucocephalus* | GCF_000737465.1 | blood sample | Illumina HiSeq 2000 | The Bald Eagle Consortium |
| *Hieraaetus wahlbergi* | GCA_035590315.1 | toe pad | Illumina HiSeqX | Iridian Genomes |
| *Nisaetus alboniger* | GCA_025447895.1 | muscle | Illumina HiSeqX | Iridian Genomes |
| *Nisaetus nipalensis* | GCA_012487455.1 | skin | Illumina HiSeqX | Environmental Genomics Office, Center for Environmental Biology and Ecosystem Studies, National Institute for Environmental Studies |
| *Pithecophaga jeffeyi* | GCA_025728025.1 | toe pad | Illumina HiSeqX | Iridian Genomes |
| *Spizaetus ornatus* | GCA_026213075.1 | muscle | Illumina HiSeqX | Iridian Genomes |
| *Spizaetus tyrannus* | GCA_013399215.1 | muscle | Illumina HiSeq 4000 | B10K Consortium |
